## Supplement for "*In silico* candidate variant and gene identification using inbred mouse strains" (from another version of this preprint: version 1, 2020-09-04)

Munz *et al.*

| Strain | Low confidence | Missing | Multi-allelic | Homozygous |
|---|---|---|---|---|
| 129P2/OlaHsd | 2759201 | 464016 | 0 | 71650637 |
| 129S1/SvImJ | 2104725 | 485850 | 987 | 72282292 |
| 129S5SvEvBrd | 6291207 | 782468 | 726 | 67799453 |
| AKR/J | 2127540 | 501461 | 5127 | 72239726 |
| A/J | 2000508 | 475908 | 5730 | 72391708 |
| BALB/cJ | 1902476 | 480064 | 6463 | 72484851 |
| BTBR | 1803722 | 424366 | 4298 | 72641468 |
| BUB/BnJ | 3445878 | 796218 | 7377 | 70624381 |
| C3H/HeH | 3107657 | 797373 | 6311 | 70962513 |
| C3H/HeJ | 1872635 | 457199 | 7565 | 72536455 |
| C57BL/10J | 1041721 | 72063 | 564 | 73759506 |
| C57BL/6NJ | 850866 | 39246 | 61 | 73983681 |
| C57BR/cdJ | 1616447 | 308528 | 4383 | 72944496 |
| C57L/J | 1819112 | 343481 | 4151 | 72707110 |
| C58/J | 1589594 | 288777 | 3776 | 72991707 |
| CAST/EiJ | 4195487 | 1189690 | 69504 | 69419173 |
| CBA/J | 1956792 | 547516 | 9055 | 72360491 |
| DBA/1J | 2111015 | 555761 | 9648 | 72197430 |
| DBA/2J | 2040804 | 592720 | 9552 | 72230778 |
| FVB/NJ | 2236272 | 401711 | 9975 | 72225896 |
| I/LnJ | 2247049 | 553205 | 13401 | 72060199 |
| KK/HiJ | 2200820 | 492819 | 15789 | 72164426 |
| LEWES/EiJ | 5459663 | 789646 | 26480 | 68598065 |
| LP/J | 2084208 | 521306 | 4361 | 72263979 |
| MOLF/EiJ | 5013644 | 1241831 | 102928 | 68515451 |
| NOD/ShiLtJ | 2704953 | 620641 | 12373 | 71535887 |
| NZB/B1NJ | 2025638 | 500590 | 15537 | 72332089 |
| NZO/HlLtJ | 1941079 | 451271 | 15282 | 72466222 |
| NZW/LacJ | 2042271 | 527620 | 15595 | 72288368 |
| PWK/PhJ | 4038588 | 1292408 | 131599 | 69411259 |
| RF/J | 3244259 | 751145 | 9947 | 70868503 |
| SEA/GnJ | 3158470 | 808689 | 9411 | 70897284 |
| SPRET/EiJ | 6367082 | 1801098 | 483567 | 66222107 |
| ST/bJ | 2075242 | 443681 | 12580 | 72342351 |
| WSB/EiJ | 2442184 | 673958 | 41658 | 71716054 |
| ZALENDE/EiJ | 2671902 | 997470 | 51720 | 71152762 |

Supplementary Table 1: For every strain the number of homozygous variants used in the database as well as the number of low confidence, missing and multi-allelic variants, all of which are excluded from fine mapping.

| Consequence | SO description | SO term | Display term | Impact | N | Percentage |
|---|---|---|---|---|---|---|
| 3_prime_UTR_variant | A UTR variant of the 3' UTR | SO:0001624 | 3 prime UTR variant | MODIFIER | 1060963 | 0.88 |
| 5_prime_UTR_variant | A UTR variant of the 5' UTR | SO:0001623 | 5 prime UTR variant | MODIFIER | 476953 | 0.39 |
| coding_sequence_variant | A sequence variant that changes the coding sequence | SO:0001580 | Coding sequence variant | MODIFIER | 276 | 0.00 |
| downstream_gene_variant | A sequence variant located 3' of a gene | SO:0001632 | Downstream gene variant | MODIFIER | 12275875 | 10.15 |
| incomplete_terminal_codon_variant | A sequence variant where at least one base of the final codon of an incompletely annotated transcript is changed | SO:0001626 | Incomplete terminal codon variant | LOW | 147 | 0.00 |
| intergenic_variant | A sequence variant located in the intergenic region, between genes | SO:0001628 | Intergenic variant | MODIFIER | 28350706 | 23.44 |
| intron_variant | A transcript variant occurring within an intron | SO:0001627 | Intron variant | MODIFIER | 35716638 | 29.54 |
| mature_miRNA_variant | A transcript variant located with the sequence of the mature miRNA | SO:0001620 | Mature miRNA variant | MODIFIER | 800 | 0.00 |
| missense_variant | A sequence variant, that changes one or more bases, resulting in a different amino acid sequence but where the length is preserved | SO:0001583 | Missense variant | MODERATE | 247063 | 0.20 |
| NMD_transcript_variant | A variant in a transcript that is the target of NMD | SO:0001621 | NMD transcript variant | MODIFIER | 5463950 | 4.52 |
| non_coding_transcript_exon_variant | A sequence variant that changes non-coding exon sequence in a non-coding transcript | SO:0001792 | Non coding transcript exon variant | MODIFIER | 2662672 | 2.20 |
| non_coding_transcript_variant | A transcript variant of a non coding RNA gene | SO:0001619 | Non coding transcript variant | MODIFIER | 22391917 | 18.52 |
| splice_acceptor_variant | A splice variant that changes the 2 base region at the 3' end of an intron | SO:0001574 | Splice acceptor variant | HIGH | 3883 | 0.00 |
| splice_donor_variant | A splice variant that changes the 2 base region at the 5' end of an intron | SO:0001575 | Splice donor variant | HIGH | 5834 | 0.00 |
| splice_region_variant | A sequence variant in which a change has occurred within the region of the splice site, either within 1-3 bases of the exon or 3-8 bases of the intron | SO:0001630 | Splice region variant | LOW | 94706 | 0.08 |
| start_lost | A codon variant that changes at least one base of the canonical start codon | SO:0002012 | Start lost | HIGH | 866 | 0.00 |
| stop_gained | A sequence variant whereby at least one base of a codon is changed, resulting in a premature stop codon, leading to a shortened transcript | SO:0001587 | Stop gained | HIGH | 3840 | 0.00 |
| stop_lost | A sequence variant where at least one base of the terminator codon (stop) is changed, resulting in an elongated transcript | SO:0001578 | Stop lost | HIGH | 841 | 0.00 |
| stop_retained_variant | A sequence variant where at least one base in the terminator codon is changed, but the terminator remains | SO:0001567 | Stop retained variant | LOW | 499 | 0.00 |
| synonymous_variant | A sequence variant where there is no resulting change to the encoded amino acid | SO:0001819 | Synonymous variant | LOW | 352392 | 0.29 |
| upstream_gene_variant | A sequence variant located 5' of a gene | SO:0001631 | Upstream gene variant | MODIFIER | 11817035 | 9.77 |

Supplementary Table 2: Variant consequences and their impact as well as the percentage of variant annotations for every consequence. Overall, the 74,480,058 variants have been annotated with 120,927,856 consequences, so on average every variant has more than one consequence. This table is taken from `https://m.ensembl.org/info/genome/variation/prediction/predicted_data.html`.
